# Stimulator of interferon genes is required for Toll-Like Receptor-8 induced interferon response

**DOI:** 10.1101/2023.05.15.540812

**Authors:** K. García-Martínez, J. Chen, J. Jones, A. Woo, A. Aucapina, I. Brito, CA Leifer

**Affiliations:** Department of Microbiology and Immunology, College of Veterinary Medicine, Cornell University, Ithaca NY 14853

**Keywords:** Innate immunity, NF-kB, IRF, Autoimmunity, Signaling, Nucleic Acids

## Abstract

The innate immune system is equipped with multiple receptors to detect microbial nucleic acids and induce type I interferon (IFN) to restrict viral replication. When dysregulated these receptor pathways induce inflammation in response to host nucleic acids and promote development and persistence of autoimmune diseases like Systemic Lupus Erythematosus (SLE). IFN production is regulated by the Interferon Regulatory Factor (IRF) transcription factor family of proteins that function downstream of several innate immune receptors such as Toll-like receptors (TLRs) and Stimulator of Interferon Genes (STING). Although both TLRs and STING activate the same downstream molecules, the pathway by which TLRs and STING activate IFN response are thought to be independent. Here we show that STING plays a previously undescribed role in human TLR8 signaling. Stimulation with the TLR8 ligands induced IFN secretion in primary human monocytes, and inhibition of STING reduced IFN secretion from primary monocytes from 8 healthy donors. We demonstrate that TLR8-induced IRF activity was reduced by STING inhibitors. Moreover, TLR8-induced IRF activity was blocked by inhibition or loss of IKKε, but not TBK1. Bulk RNA transcriptomic analysis supported a model where TLR8 induces transcriptional responses associated with SLE that can be downregulated by inhibition of STING. These data demonstrate that STING is required for full TLR8-to-IRF signaling and provide evidence for a new framework of crosstalk between cytosolic and endosomal innate immune receptors, which could be leveraged to treat IFN driven autoimmune diseases.

**Background:** High levels of type I interferon (IFN) is characteristic of multiple autoimmune diseases, and while TLR8 is associated with autoimmune disease and IFN production, the mechanisms of TLR8-induced IFN production are not fully understood.

**Results:** STING is phosphorylated following TLR8 signaling, which is selectively required for the IRF arm of TLR8 signaling and for TLR8-induced IFN production in primary human monocytes.

**Conclusion:** STING plays a previously unappreciated role in TLR8-induced IFN production

**Significance:** Nucleic acid-sensing TLRs contribute to development and progression of autoimmune disease including interferonopathies, and we show a novel role for STING in TLR-induced IFN production that could be a therapeutic target.

## Introduction

Interferonopathies are autoinflammatory disorders that are characterized by high serum levels of type-I interferon (IFN) due in some cases to inappropriate activation of innate immune receptors by nucleic acids (1–10). IFN plays a pathologic role in many of these diseases, including Systemic Lupus Erythematosus (SLE) (11) where increased IFN expression is correlated with disease activity (12). Anifrolumab, an IFNAR blocking antibody, improves cutaneous manifestations and arthritis in SLE patients (13, 14); however, this therapy fails in more than 50% of patients with high IFN signatures (15). Thus, targeting components of the IFN induction pathway provides another therapeutic option for interferonopathies.

Activation of a subset detect nucleic acids (TLR7/8/9) induces IFN secretion and contributes to SLE pathogenesis (16–19). While plasmacytoid dendritic cells (pDCs) express TLR7 and TLR9 and are considered the major source of IFN in SLE (20), TLR8 is not expressed in pDCs. Yet, TLR8 mutations, and TLR8 expression levels in other cell types, is associated with autoimmune disease (21–24). Moreover, sera obtained from SLE patients activates both TLR7 and TLR8 in reporter cells (25), increased TLR8 activation is observed in peripheral blood mononuclear cells from pediatric SLE patients (26), and immune complexes from SLE patients activate TLR8 (27). Monocytes lack TLR7 and TLR9, but express TLR8 (28) and are significantly more numerous than pDCs in the blood (29). Indeed, monocytes are the main source of IFN in some mouse models of SLE (30). TLR8 signaling events are generally assumed to be the same as TLR7 and TLR9 as these TLRs respond to nucleic acids and relay their signal through the same co-adaptor protein, MyD88. Yet, TLR7 and TLR8 are respond to different ligand structures suggesting there are important differences for how TLR8 couples to its effector pathways compared to TLR7 and TLR9. TLR8 is thus a potential therapeutic target to regulate IFN levels in autoimmune interferonopathies arguing for a need to understand its molecular mechanisms of signaling.

The activation of the cyclic GMP-AMP synthase (cGAS) - stimulator of interferon genes (STING) pathway is another nucleic acid sensing pathway implicated in SLE (31) and interferonopathies, such as STING-Associated vasculopathy with onset in infancy (SAVI) (2, 32) and Aicardi–Goutières syndrome (AGS) (33). Thus, the STING pathway is a potential therapeutic target for these diseases (34). Microbial, nuclear, or mitochondrial DNA is detected by cytoplasmic cGAS, which then generates the second messenger cGAMP that subsequently binds to and activates STING (35). Activated STING multimerizes and traffics from the ER to the Golgi where it acts as a scaffold to recruit TANK-binding kinase 1 (TBK1) and interferon regulatory factors (IRFs) to ultimately induce IFN production (36). While both TLRs and STING activate TBK1, a critical kinase that activates IRFs, these pathways are considered to be independent. However, there are potential connections between TLR signaling and the cGAS/STING pathway since TLR engagement can induce mitochondrial DNA release and activation of cGAS (37).

In this study, we demonstrate that the STING signaling pathway has a previously unappreciated, and selective, crosstalk with human TLR8 that is required for TLR8 to induce IFN production in human monocytes. TLR8 stimulation induces rapid STING phosphorylation in primary human monocytes and STING inhibition significantly reduces IFN production by both the THP-1 monocyte cell line and primary monocytes. STING inhibition selectively reduces TLR8-induced IRF activity, but not NFkB activity. TLR8-induced IRF activation is independent of IFN secretion and IFN alpha receptor (IFNAR). Finally, RNA sequencing revealed that TLR8 induces many SLE-associated genes and that those disease-associated genes are significantly downregulated by STING inhibition. Collectively, these studies expand our understanding of TLR8 signaling and regulation and build a new framework for crosstalk between cytosolic and endosomal innate immune receptors that could be used to target autoimmune diseases mediated by high IFN levels.

## Results

### STING inhibition decreases TLR8-induced Type I IFN secretion

To determine whether STING is involved in TLR8 mediated IFN secretion, we treated primary monocytes from healthy donors with increasing concentrations of the STING inhibitor, H151, then stimulated with the TLR8 ligand TL8-506 (TL8). STING inhibitor pretreatment reduced TLR8-induced IFN secretion in a concentration dependent manner (Fig. 1A). This contrasted with no effect of STING inhibition on IL-6 production measured in the same supernatants (Fig. 1B). H151 decreased both IFN and IL-6 secretion from cells treated with cGAMP, a STING agonist (Fig. 1A-1B). At all of the concentrations used, H151 did not affect viability of the primary monocytes (Fig. 1C). IFN secretion by primary monocytes from 8 independent donors was induced by TLR8 stimulation and reduced in the presence of STING inhibition (Fig. 1D). STING inhibition also decreased both TLR8 and cGAMP induced IFN secretion in THP-1 cells, a monocytic cell line (Fig. 1E). However, THP-1 cells were more sensitive to toxicity with higher H151 concentrations than monocytes (Fig. 1F); therefore, we used the lowest concentration of H151 for our subsequent THP-1 experiments. We conclude that STING is required for full TLR8-induced IFN secretion.

**Figure 1:**
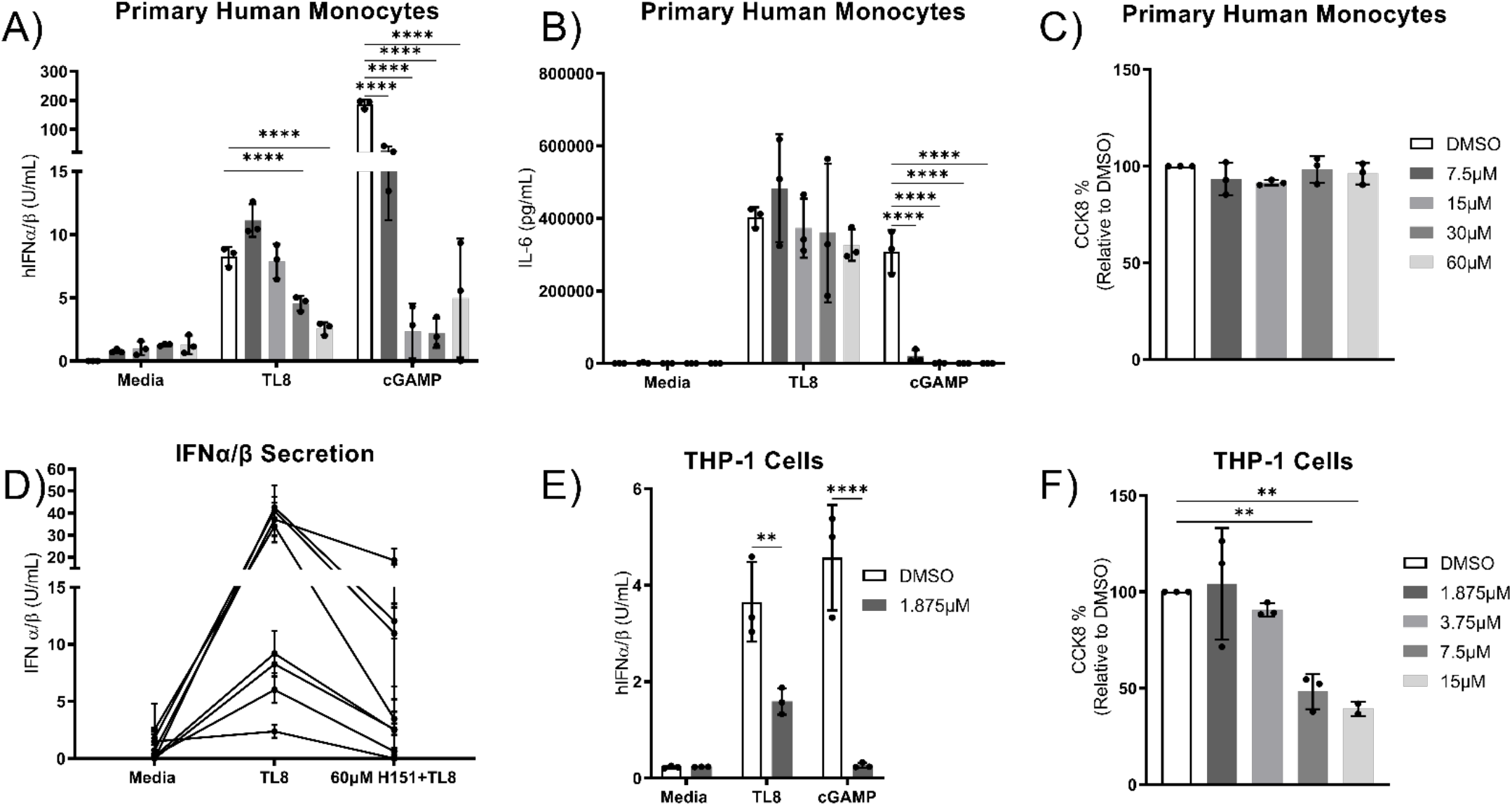
STING inhibition decreases TLR8-mediated Type I IFN secretion. Primary human monocytes and THP-1 cells were treated with vehicle or indicated concentrations H151 for 1 hour, then stimulated with TL8-506 (TLR8 ligand), or cGAMP (STING ligand) for 18 hours. Supernatants were collected to measure IFNα/β secretion using IFNα/β Reporter HEk293 cells. CCK8 activity was measured in parallel. A) Representative experiment showing IFNα/β secretion in primary human monocytes n=4.B) Representative experiment showing IL-6 secretion from primary monocytes n=4. C) Representative experiment showing CCK8 activity in primary human monocytes n=4. D) IFN α/β secretion from 8 different donors n=8. E) Representative experiment showing IFNα/β secretion in THP-1 cells n=3. F) Representative experiment showing CCK8 activity in THP-1 cells n=4. Data are presented as means ± S.D. (error bars) from a representative experiment (2way ANOVA with multiple-comparison test). ****=p<0.0001, **=p<0.01.

### STING inhibition abrogates TLR8-mediated IRF, but not NFkB, activity

The two major transcription factors activated downstream of the TLR signaling pathways are IRFs and NFkB (38–40). Our results from Figure 1 show that H151 decreases IFN, an IRF dependent cytokine, but not IL-6, a predominantly NFkB dependent cytokine. Therefore, we tested whether STING is required selectively for IRF activation. Dual Reporter THP-1 (DTHP-1) cells stably express reporters for IRF and NFkB activity, which allows measurement of both pathways simultaneously from the same cells. Knockout of STING in DTHP-1 cells eliminated both IRF and NFkB activity in response to the TLR7/8 ligand R848 (WT cells do not express TLR7 since they don’t respond to TLR7-specific imiquimod or loxoribine) and eliminated IRF activity in response to the STING ligand cGAMP without affecting IFN induced IRF activity (Fig. S1). Lack of TLR8 mRNA expression in STINGKO cells, which we observed in our RNA sequencing data, likely accounts for this lack of TLR8 signaling. Therefore, we treated DTHP-1 cells with vehicle control or two different STING inhibitors with different mechanisms of action prior to TLR8 stimulation to test the role of STING in TLR8 signaling. H151 covalently binds to STING and irreversibly blocks STING palmitoylation and clustering (41), while SN-011 binds to the cyclic dinucleotide-binding pocket of STING, blocking cGAMP binding (42). Both STING inhibitors significantly reduced TL8-induced IRF activity. As expected, stimulation with cGAMP and IFNβ also increased IRF activity and cGAMP-induced IRF activity was completely abrogated by both STING inhibitors while there was little to no effect on IFNβ-induced IRF activity (Fig 2.A). Moreover, H151 inhibition did not decrease TLR8-induced NFkB activity, yet SN-011 did reduce TLR8 induced NFkB activity (Fig. 2B) without reducing cell viability (Fig. S2A).

**Figure 2:**
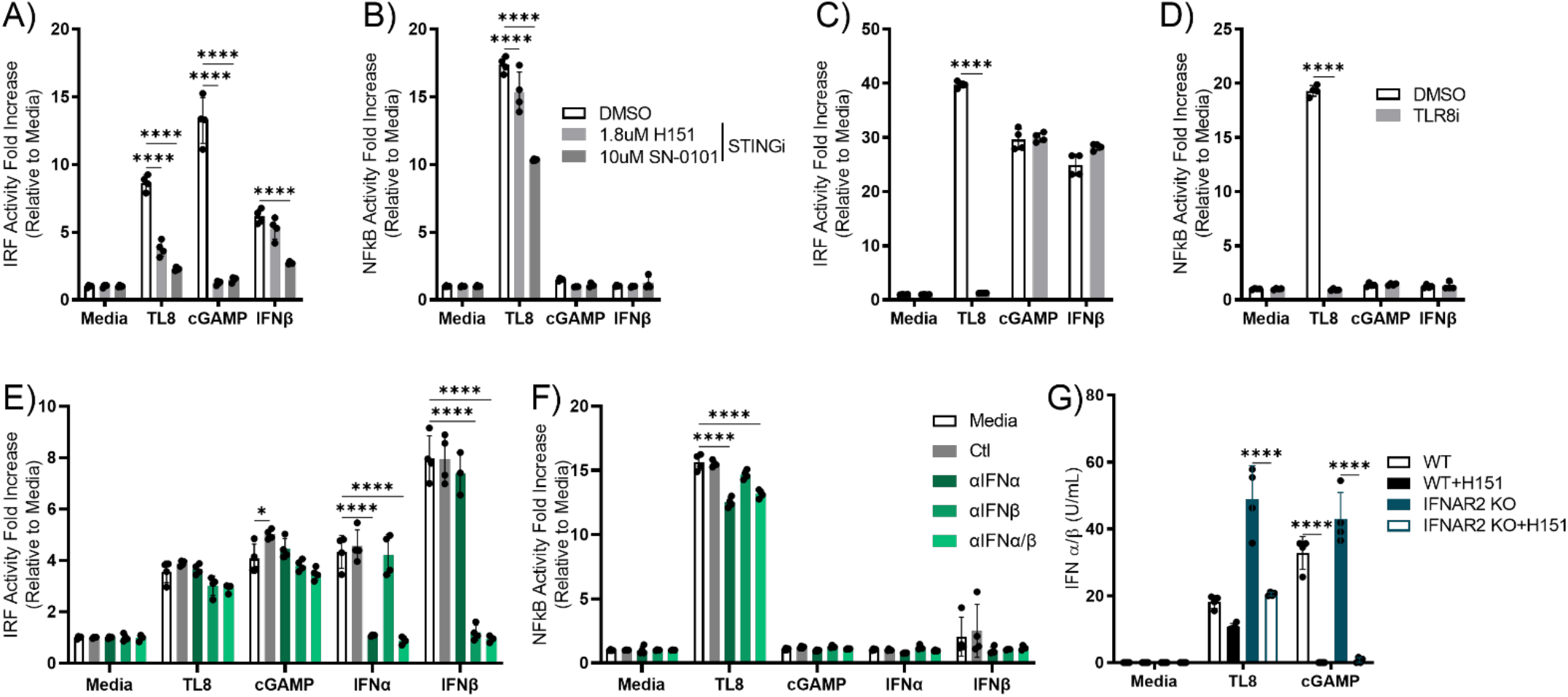
STING inhibition abrogates TLR8-mediated IRF activity, but not NFkβ. DTHP-1 cells were treated with vehicle, 1.8μM H151, 10μM SN-011 for 1 hour, then stimulated with TL8 for 18 hours cGAMP and hIFNβ were used as positive controls for STING dependent and independent IRF activation. IRF activity was measured using Lucia Luciferase activity and NFkβ activity was measured using Secreted Alkaline Phosphatase (SEAP) activity. A) Representative experiment showing IRF activity n=6. B) Representative experiment showing NFkB activity n=6. DTHP-1 cells were treated with vehicle, or 1μM CUCPT9a for 1 hr, then stimulated with TL8, cGAMP, or hIFNβ for 18 hours. C) Representative experiment showing IRF activity n=3 D) Representative experiment showing NFkβ activity n=3. DTHP-1 cells were stimulated with TL8, cGAMP, IFNα, or IFNβ in the presence of media, isotype control antibodies, 1:7,000 anti-IFNα, 1:1,000 anti-IFNβ, or a combination of anti-IFNα/β neutralizing antibodies for 18 hours E) Representative experiment showing IRF activity n=4. F) Representative experiment showing NFkβ activity n=4. WT or IFNAR2KO cells were inhibited with 1.875 μM H151 or DMSO for 1hr, then stimulated with 10 μg/mL TL8 or cGAMP for 18 hours. Supernatants were collected and assayed for IFN secretion. G) Representative experiment showing IFN secretion n=3. Data are presented as means ± S.D. (error bars) from a representative experiment (2way ANOVA with multiple-comparison test). ****=p<0.0001, **=p<0.01, *=p<0.05.

Since TL8 is a synthetic benzoazepine compound (43, 44), we next tested whether TL8 might have TLR8-independent activity. We treated the cells with CUCPt9a, a TLR8 specific inhibitor, which reduced IRF and NFkB activity to baseline (Fig. 2C-D). CUCPt9a did not inhibit STING mediated IRF activity by cGAMP, nor IRF activity mediated by the interferon-α/β receptor (IFNAR) through IFNβ (Fig. 2C). CUCPta selectively abrogated TL8 induced IRF activity and NFkB activity and not LPS mediated activity (Fig. S2C-D) without affecting cell viability (Fig. S2B).

Since IFN is secreted in response to TLR8 stimulation and IFN can also induce IRF activity, we next tested if TLR8-induced IRF activity was indirectly induced through a positive feedback loop (45). DTHP-1 cells were stimulated with TL8, cGAMP, IFNα, or IFNβ in the presence of isotype control antibodies or IFN neutralizing antibodies. Anti-IFNα neutralized IFNα-induced IRF activity, and anti-IFNβ neutralized IFNβ-induced IRF activity (Fig. 2E). The mix of anti-IFNα/β neutralized both IFNα and IFNβ-induced IRF activity (Fig 2E). In contrast, neither neutralizing antibody alone, or the combination, had any effect on TL8-induced or cGAMP-induced IRF activity (Fig. 2E), while there was a slight reduction of TL8-induced NFkB activity with anti-IFNα and the mix (Fig. 2F). Importantly, in IFNAR2KO cells TL8 still induced IFN secretion, which is reduced upon STING inhibition (Fig. 2G). These data support a direct effect of STING in TL8-induced IRF activity independent of any IFN/IFNAR autocrine feedback loop.

### TL8 induces STING phosphorylation and requires IKKε, but not TBK1, to activate IRFs

A key event in STING activation is its phosphorylation at amino acid Serine 366 by the kinase TBK1 (46). Monocytes stimulated with three different TLR8 ligands, TL8, ssPolyU, and ssRNA40, had detectable Serine 366-phosphorylated STING within 30 minutes (Fig. 3A). Stimulation with TL8 also induced TBK1 phosphorylation.(Fig. 3B). IFN neutralizing antibodies did not decrease TL8-induced STING phosphorylation (Fig 3C). Moreover, IFNα or IFNβ did not induce STING phosphorylation, but did induce STAT1 phosphorylation, which was inhibited by the neutralizing antibodies (Fig. 3C). Importantly, TL8 stimulation did not induce STAT1 phosphorylation (Fig. 3C). Together, these data support a model where TL8 induces rapid and direct STING phosphorylation independent of IFN production or signaling.

**Figure 3:**
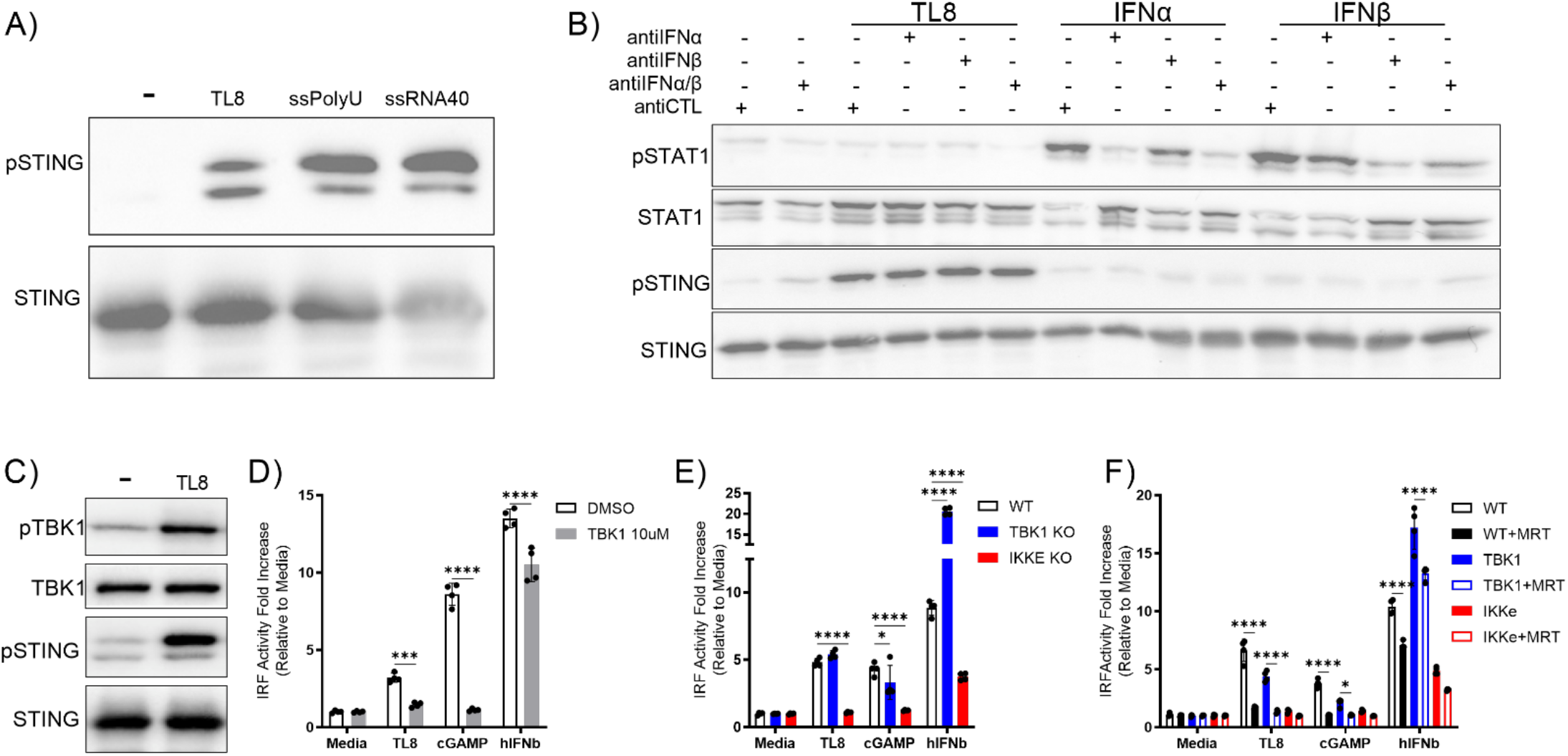
TL8 induces STING phosphorylation and requires IKKε, but not TBK1. A) Human monocytes were stimulated with 10ug/mL TL8, 10ug/mL ssPolyU, or 10ug/mL ssRNA40 for 30 minutes. Cell lysates were resolved by SDS-PAGE and immunoblotted for phosphorylated STING (p40kDa), then stripped and re-probed for total STING (p35kDa) blot from representative experiment n=3. B) Human monocytes were co-treated with control antibody, 1:7,000 anti-IFNα, or 1:1000 anti-IFNβ antibodies and 10ug/mL TL8, 100U/mL IFNα, and 100U/mL IFNβ for 30 minutes. Cell lysates were resolved by SDS-PAGE and immunoblotted for phosphorylated STAT1 (85kDa) and phosphorylated STING (40kDa, then stripped and re-probed for total STAT1 (84kDA) and total STING (p35kDa) blot from representative n=3.C) THP-1 cells were stimulated with media or 10 μg/mL TL8. Cell lysates were resolved by SDS-PAGE and immunoblotted for phosphorylated STING (p40kDa) and phosphorylated TBK1 (75kDa), then stripped and re-probed for total STING (p35kDa) and total TBK1 (75kDa) blot from representative experiment n=3. WT DTHP-1 cells were treated with vehicle, 10μM MRT67307 for 1hr then stimulated with TL8, cGAMP, or hIFNβ for 18 hours. D) Representative experiment showing IRF activity, n=6. TBK1 KO, or IKKε KO DTHP-1 cells were stimulated with TL8, cGAMP, or hIFNβ for 18 hours. E) Representative experiment showing IRF activity, n=6. WT, TBK1 KO, or IKKε KO DTHP-1 cells were treated with vehicle or 10μM MRT67307 for 1hr then stimulated with TL8, cGAMP, or hIFNβ for 18 hours. F) Representative experiment showing IRF activity, n=3. Data are presented as means ± S.D. (error bars) from a representative experiment (2way ANOVA with multiple-comparison test). ****=p<0.0001, ***=p<0.001, *=p<0.05.

MRT67307 (MRT), a TBK1/IKKε inhibitor (47), abrogated TLR8-induced IRF activity (Fig. 3D), but not NFkB activity (Fig. S3A). MRT also reduced cGAMP-induced IRF activity and slightly reduced IFN induced IRF activity (Fig. 3D). Surprisingly, TL8 induced IRF activity was similar in TBK1 deficient cells compared to WT cells (Fig. 3E). Yet, TL8-induced IRF activity was completely abrogated in the absence of IKKε (Fig. 3E). NFkB activity was slightly reduced in both TBK1 and IKKε deficient cells (Fig. S3B). Inhibition with MRT further reduced TL8-induced IRF activity in TBK1 KO cells (Fig. 3F) without reducing NFkB activity (Fig. S3C). Thus, TLR8 stimulation induces STING phosphorylation and TLR8-induced IRF activity depends on IKKε and not TBK1.

### STING inhibition decreases TLR8-induced autoimmune disease-associated genes

Since STING is a potential therapeutic target for certain autoimmune interferonopathies (34), we next tested if TLR8 induces a set of genes via STING that may play a role in pathogenesis. To identify STING-dependent TLR8-induced genes, we performed transcriptomic analysis on TLR8-stimulated THP-1 cells with or without STING inhibition. TLR8 stimulation upregulated 775 genes and 85 were significantly decreased upon STING inhibition by H151 (Fig. 4A). We performed GO Biological Processes pathway and Molecular Signature Hallmark analyses and identified multiple IFN related pathways that were induced by TL8 stimulation (Fig. 4B-C). Pathway analysis using DisGeNET showed TLR8-associated signatures included viral infection, inflammation, cancer, and autoimmune diseases (Fig. S4). Narrowing the analysis to interferon-associated and immunopathology annotations we analyzed the 30 most significantly associated diseases. Our analysis revealed an over representation of Lupus diseases and the presence of ‘interferonopathy’ described diseases like Sjorgen’s Syndrome (Fig. 4D). A Gene Set Enrichment Analysis (GSEA) showed that TLR8 stimulated, compared to unstimulated cells, had a high positive Normalized Enrichment Score (NES) for SLE (NES 4.46). Importantly, the SLE enrichment score was highly downregulated when STING was inhibited by H151 in TL8 stimulated cells compared to uninhibited cells (NES -2.07) (Fig. 4E-F). Together, we conclude that TLR8 induces a set of STING-dependent genes that are associated with SLE. Thus, STING inhibition is a potential therapeutic target to reduce nucleic acid sensing TLR-induced IFN production that contributes to the pathology in several autoimmune diseases.

**Figure 4:**
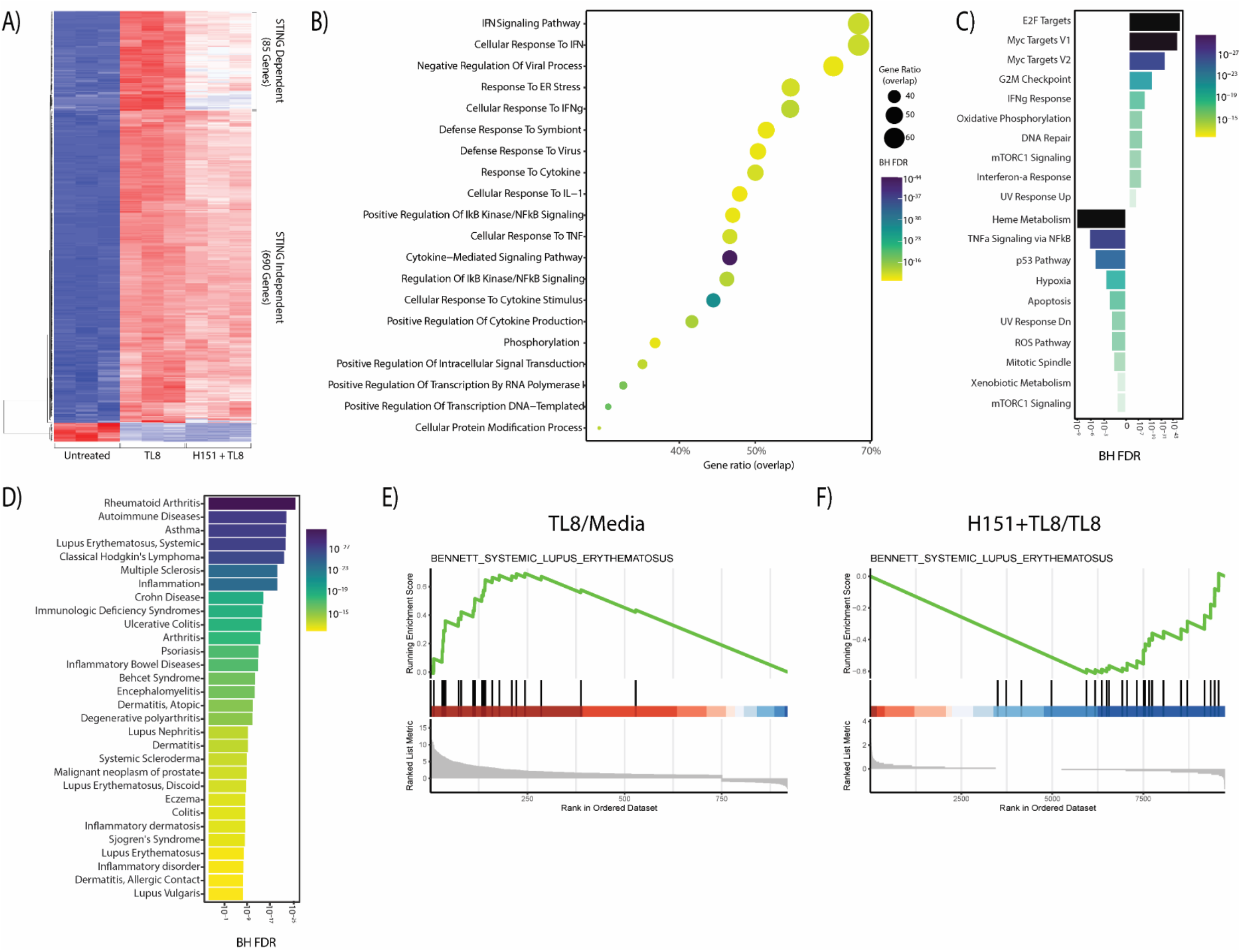
STING inhibition decreases TLR8 induced genes associated with autoimmune diseases. THP-1 cells were inhibited with 500ng/mL H151 or DMSO for one hour, then treated with media or 10μg/mL TL8 for 8 hours in triplicates per condition. RNA was isolated by trizol and bulk RNA sequencing was performed. A) Heatmap of RNA-sequencing analysis showing genes upregulated by TL8 stimulation, data shown in triplicates. B) GO pathways induced by TLR8 stimulation. C) Bar plot depicting the top 10 upregulated and downregulated gene sets for H151TL8 stimulated cells vs TL8 stimulated cells. D) Bar plot depicting 30 significantly upregulated interferon-associated and immunopathology annotations according to DisGeNET for TL8 stimulated cells, plotted in descending according to corrected p-values. E) Results from Gene Set Enrichment Analysis (GSEA) comparison between TL8 stimulated and unstimulated THP-1 cells for SLE associated genes and F) GSEA comparison between H151 inhibited and uninhibited cells stimulated with TL8 for SLE associated genes.

**Figure 5:**
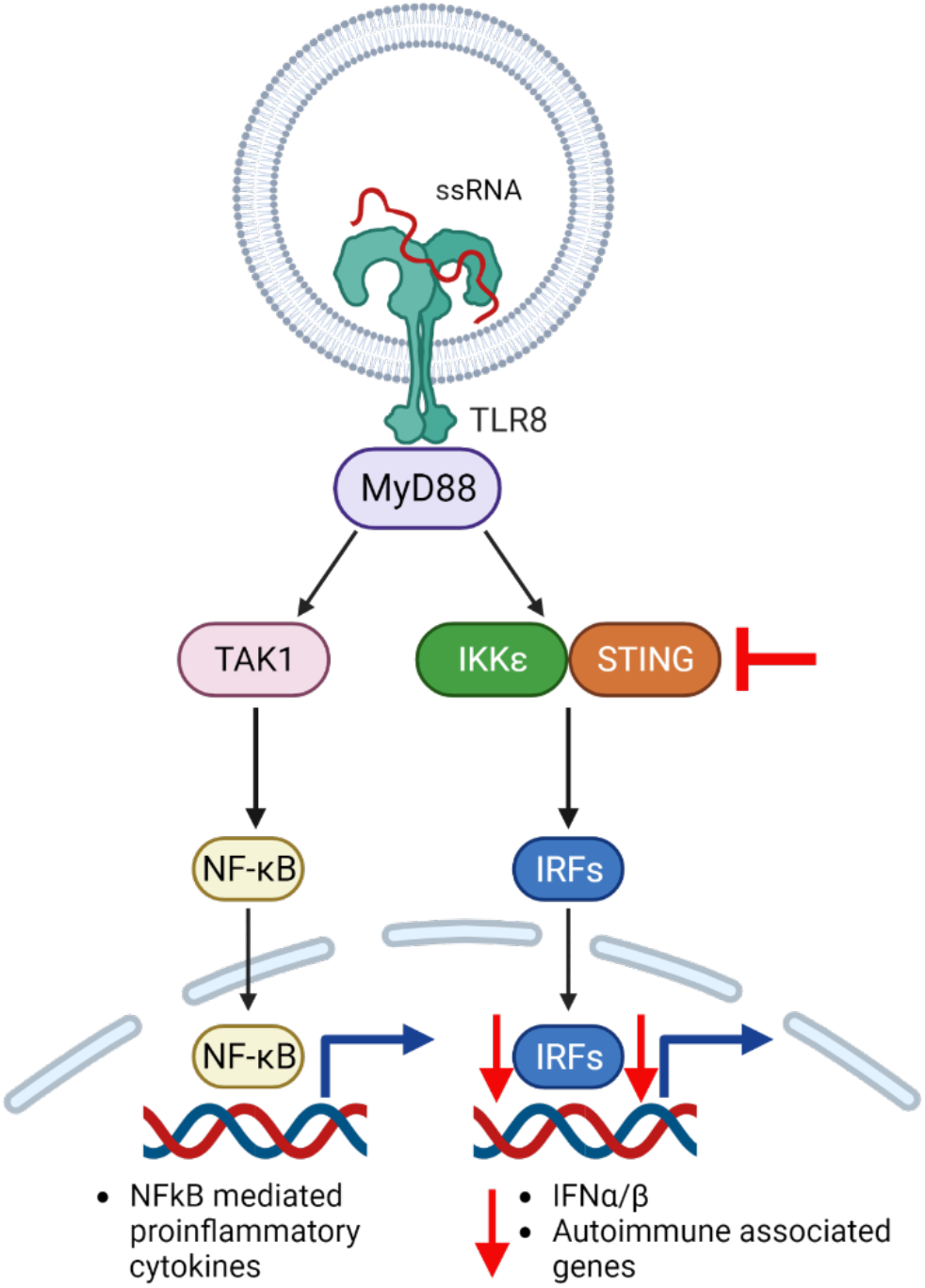
Model for role of STING in TLR8-induced IRF signaling. TLR8 is activated upon ssRNA binding to receptor and transduces the signal through MyD88. This leads to a NFkB/IRF signal bifurcation. NFkB signaling is induced via TAK1 and results in NFkB mediated proinflammatory cytokines, which is unaffected by STING inhibition. TLR8 activation also leads to IRF signaling through MyD88. Upon activation, MyD88 recruits IKKε and STING, with lead to activation of IRFs and upregulation of IFNα/β genes and autoimmune associated genes. STING inhibition before TLR8 activation selectively reduces IRF mediated signaling which results in reduction of IFNα/β genes and autoimmune associated genes.

## Discussion

IFN production induced by TLR2, TLR3, and TLR4, is initiated by the adapter protein TIR-domain-containing adapter-inducing interferon-β (TRIF), linking TRAF3 to TBK1 resulting in IRF phosphorylation and subsequent IFN induction. Yet, the endosomal TLRs, TLR7, TLR8, and TLR9, which exclusively use MyD88 as an adapter protein also lead to IFN production. The link between these endosomal sensors is less well understood mechanism. It is known that upon activation in pDCs, TLR7 and TLR9 bind MyD88 which subsequently recruits IL-1 receptor-associated kinase (IRAK) to TLRs. IRAK binds to TNF receptor-associated factor-6 (TRAF6) and this complex activates transforming growth factor-β-activated protein kinase 1 (TAK1) leading to NF-κB activation. IRAK/TRAF6 also activates IRF7, which is the main regulator of IFN production in pDCs. While pDCs express TLR7 and TLR9 (20), they do not express TLR8, which is expressed highly in monocytes (21). Differences in signaling pathway activation have been observed between TLR7 and TLR8 between immune cells (25). Thus, it is important to understand TLR8 signaling in monocytes, which are an underappreciated source of IFN in autoimmune diseases (30, 48–50). Our study shows that TLR8 induces IFN production in a STING dependent manner in monocytes and supports a new model for selective regulation of TLR8 mediated IFN in monocytes that could be exploited for IFN modulation in autoimmune diseases.

TLR activation may indirectly induces IFN production by increasing mitochondrial permeability, which in turn releases mtDNA into the cytoplasm that binds cGAS. cGAS produces the second messenger cGAMP, which binds and activates STING (51, 52). This pathway can result in IFN secretion that can suppress viral replication, but can also contribute to autoimmune pathology (53). Our studies support an alternative model where TLR8 activated STING and IRF activity independently of cGAS since cGAS KO cells stimulated with TL8 still induced IRF and NFkB activity, as well as TL8 dependent IFN secretion (Fig. S5A-C).

More recent studies describe a direct interaction between MyD88 dependent TLRs and IFN response machinery. Tan and Kagan found that in the absence of TRIF, MyD88 expressing cells can induce TBK1 phosphorylation upon TLR2, TLR4, and TLR7/8 stimulation through the supramolecular organizing center called the myddosome. In the myddosome, MyD88 and TBK1 physically interact and induce MyD88 dependent IFN response (54). TBK1 activation by the myddosome increased glycolysis, without activating NF-kB, showing a selectivity for the IRF pathway. Recently, the proteins TASL and SCL15A4 were also shown to induce IRF, specifically IRF5, downstream of TLR7 and TLR9; however, both of these proteins were involved in both the IRF and NFkB pathways and could not account for our observation of selective regulation of the IRF pathway (55).

Our data support a direct role for STING in TLR8-induced IFN production. We found that STING inhibition, before TLR8 stimulation, significantly reduced IFN secretion in primary human monocytes from healthy donors and THP-1 cells without significantly reducing TLR8-induced IL-6 secretion (Fig. 1). We also observed that STING inhibition selectively decreased TLR8 mediated IRF activity to almost to basal levels (Fig. 2A). TLR8 induced IRF activity was independent of an IFN positive feedback loop (Fig 2D) that has previously been described for TLR2 (56). In the cGAS-STING pathway, STING functions as a scaffolding protein that promotes the TBK1-IRF3 interaction (57), our data suggests that STING may also function as a scaffold in the TLR8-IRF pathway.

TLR8 stimulation induced STING phosphorylation within 30 minutes, which was also independent of an IFN positive feedback loop (Fig. 3A-B). These data support a direct role for STING in TLR8 induced IRF activity. Previous reports identify TBK1 as the kinase responsible for MyD88 dependent IRF activity (54, 58), however these studies use TBK1/IKKε double knockouts or inhibitors that inhibit both TBK1 and IKKε activity. Using individual knockouts for TBK1 and IKKε, our data demonstrate that TLR8 induced IRF activity is dependent on IKKε but not TBK1. In fact, IRF activity was induced similarly in WT and TBK1KO cells, and TBK1/IKKε inhibitor reduced IRF activity to baseline in both WT and TBK1KO cells (Fig. 3C-F).

The cGAS-STING pathways is a potential therapeutic target for many inflammatory and autoimmune diseases (34). For example, STING deficiency improves pathology in an AGS mouse model (59). STING knockout in 129/Fcgr2b^-/-^ mice, a mouse line that develops SLE-like disease, rescues SLE symptoms and increases survival. However, knocking out STING inhibited DC maturation and pDC differentiation, so it is unclear if it there were additional effects that could be attributed to reduced monocyte signaling and subsequent reduced IFN production (31). cGAS and STING knockdown in fibroblast-like synoviocytes from rheumatoid arthritis patients resulted in lower secretion of proinflammatory cytokines like TNFα, IL-1β, and IFNβ compared to wild type synoviocytes (60). STING activation in this model was attributed to detection of improperly cleared dsDNA in the cytoplasm by cGAS; however, nucleic acids can also activate endosomal TLRs and many of these diseases are associated with TLR activation by nucleic acids (61–64). Our studies now show that STING regulates a subset of TLR8-induced genes associated with SLE and that blocking STING may be therapeutically useful for reducing TLR8-dependent signaling in autoimmune disease (Fig. 4A-D). Inhibiting STING selectively reduced TLR8-induced IFN while leaving the NF-kB pathway mostly intact, which may provide a therapeutic mechanism to retain critical antimicrobial monocyte/macrophage activities while reducing IFN production and IFN-mediated pathology. Our data also advance our knowledge on TLR8 signaling pathways, which are poorly studied. We provide evidence for a new framework of crosstalk between cytosolic and endosomal innate immune receptors, which could be leveraged to treat IFN driven autoimmune diseases.

## Methods

### Cell culture

The human acute monocytic leukemia cell line containing dual IRF and NFkB activity reporters, THP1-Dual cells (InvivoGen) were cultured and maintained in RPMI 1640 (Corning) supplemented with 10% FBS, 50 U ml−1 penicillin, 50 μg ml−1 streptomycin, 100 μg ml−1 normocin, and 2 mM L-glutamine.100 μg ml−1 of zeocin and 10 μg/ml−1 of blasticidin were added to growth medium every other passage to retain the dual reporters. WT and STING KO cells were purchased from Invivogen and cGAS and IFI16 THP1-Dual cells were created using the CRISPR Cas9 system (see below). Human Primary monocytes were obtained from University of Nebraska Medical Center and cultured in Dulbecco’s modified Eagle’s medium (DMEM) (Corning#10-017-CM), 10% human serum (SeraCare #1830-0003), 2mM L-glutamine (Corning #25-005-CI 200mM), 10mM HEPES (Gibco 15630-080 1M), 1mM Sodium Pyruvate (Corning #25-000-CIR 100mM), 50 IU Pen/Strep (Corning #30-001-CI 5,000IU). Upon delivery, 50ug/mL gentamicin were added to the primary monocytes for 15 minutes at 4C to kill any potential contamination from shipping and handling. Then cells were washed and cultured in complete DMEM for all experiments.

### Cell Stimulations

Stimulations for dual reporter and ELISA assays, THP-1 Dual cells were plated 1x10^5^ cells in 180uL per well in 96 well plate. For inhibitor studies, cells were inhibited with the STING inhibitor H151 with the indicated concentrations, 1 hr before TLR and cytosolic sensor stimulations. Cells stimulated with TLR ligands 1ug/mL Pam3CSK4 (TLR1/2), 1ug/mL LPS (TLR4), and 10ug/mL TL8-506 (TLR8). 10ug/mL of cGAMP were used to stimulate STING and 1ug/mL VACV70 using Lyovec as a transfecting reagent were used to stimulate cGAS. All stimulations were done for 18 hours. Plates were then spun at 2,000xg for 2 minutes and supernatants were collected for Lucia Luciferase, SEAP, and ELISA assays.

Primary human monocytes were seeded at 1.5x10^5^ cells in 180uL per well. For inhibitor studies, cells were inhibited with the STING inhibitor H151 with the indicated concentrations, 1 hr before TLR and cytosolic stimulations. Cells stimulated with TLR ligands 1ug/mL LPS (TLR4), and 10ug/mL TL8 (TLR8). 10ug/mL of cGAMP were used to stimulate STING. All stimulations were done for 18 hours. Plates were then spun at 2,000xg for 2 minutes and supernatants were collected for ELISA assays.

### Western Blot

Human monocytes were stimulated with 10ug/mL TL8 (Invivogen, tlrl-tl8506), 10ug/mL ssPolyU(Invivogen, tlrl-lpu) 10ug/mL ssRNA40 (Invivogen, tlrl-lrna40), 100U/mL IFNα (BEI, No. Ga23-902-530), or 100U/mL IFNβ (BEI, No. Gb23-902-531) for 30 minutes. Stimulated cells were washed with ice-cold PBS and lysed for direct immunoblotting with 13 SDS-PAGE–reduced sample buffer (62.5 mM Tris [pH 6.8], 12.5% glycerol, 1% SDS, 0.005% bromophenol blue, 1.7% 2-ME) or for immunoprecipitation (IP) or coimmunoprecipitation with lysis buffer (50 mM Tris-Cl [pH 7.4], 150 mM NaCl, 10% [w/v] glycerol, 1 mM EDTA, and protease inhibitors). Lysates were incubated at 95°C for 5 min prior to resolving by 10% SDS-PAGE. Proteins were transferred to nitrocellulose membranes and immunoblotted with the indicated Abs. Membranes were incubated with a SuperSignal West Pico chemiluminescence Western blotting detection system (Thermo Scientific) imaged using Biorad ChemiDoc. For IP experiments, total protein was determined in clarified lysates using the BCA protein assay (Bio-Rad), and 5 mg of indicated Ab was used for IP. For coimmunoprecipitations, one SDS-PAGE gel was run and transferred to nitrocellulose. The blots were sequentially probed with the indicated Abs.

### Lucia luciferase and SEAP assays

QUANTI-Luc luciferase reagent (InvivoGen) was used to measure IRF activity through Luciferase luminescence following the manufacturer’s protocol. In brief, 10 μl of cell culture supernatant was transferred to 96-well white opaque plate and luminescence was read using a Luminometer with an autoinjector and 50 μl of luciferase reagent. NfkB activity was measured in the same supernatants using 20uL of supernatant with180uL QUANTI-Blue SEAP detection reagent (InvivoGen). The samples were then incubated at 37 °C for 1 h and absorbance was measured at 640 nm. Data was represented as fold increase compared to media control for each cell line, for example WT Stimulated/WT media and STINGKO Stimulated/STING KO media.

### Cytokine Detection Assays

Cells were stimulated as indicated above. IL-6 secretion was detected using BioLegend IL-6 ELISA kit. Type I IFN secretion was detected Hek-Blue IFNa/b Cells (Invivogen). Briefly, 20 uL of supernatant from treated cells were transferred to a flat-bottom 96-well plate. 50,000 HEK-Blue IFN-α/β cells in 180 μl were added to each sample and incubated overnight at 37 °C in 5% CO2. The next morning, 20 μl of induced HEK-Blue IFN-α/β cells supernatant were mixed with 180 μl of QUANTI-Blue (invivogen) and incubated for 37 °C for 1hr. Plate was then read using a spectrophotometer 650nm.

### Stable Gene knockout through CRISPR Cas9

All guide RNAs for this study were designed using https://crisprgold.mdc-berlin.de/index.php and synthesized by Integrated DNA Technologies. sgRNAs used were a following: cGAS forward sgRNA primer: CACCGCGGCCCCCATTCTCGTACGG, cGAS reverse sgRNA primer: AAACCCGTACGAGAATGGGGGCCGC. TBK1 forward sgRNA primer CACCGTCCCCGTCTAAATCATAACG, TBK1 reverse sgRNA primer AAACCGTTATGATTTAGACGGGGA. Dual THP-1 cells were generated using lentiviral transduction. To generate lentiviral particles, 10ug of lentiviral vector (lentiCRISPR v2 (Plasmid #52961, Addgene) containin g sgRNA constructs, 2.5ug of packaging plasmid pMD2.G (Plasmid #12259, Addgene), and 7.5ug of envelope plasmid psPAX2 (Plasmid #12260, Addgene) were diluted in 250uL Opti-MEM (Gibco). The components were mixed with 30 μl FuGene (Promega) in 250 Opti-MEM and incubated for 5 min at RT. The solutions were then mixed and incubated for 20 mins at RT. Solution was added drop by drop to Hek293T cells in complete DMEM at 40% confluency. Media was replaced the 16 hours after transfection. Then supernatant was collected after 48 hours and media was replaced. Supernatant was collected after an additional 24 hours. Cells were spun at 400xg for 5 mins to remove debris after each collection. Then supernatant was spun at 2,000xg for an additional 20mins before ultracentrifugation at 100,000xg to concentrate virus in 1mL. 100uL was added to 250,000 THP-1 cells with 8ug/mL polybrene. After 48 hours cells were selected using 0.8ug/mL puromycin selection. Knock out efficiency was analyzed by immunoblotting cell lysates for TBK1, cGAS, IFI16.

### RNA sequencing

1x10^6^ cells THP-1 cells were exposed to media or H151 for 1 hour then stimulated with LPS, 10ug/mL TL8-506 and 10ug/mL of cGAMP for 8 hours. RNA was isolated using Trizol LS (Thermofisher). RNA integrity was confirmed by Fragment Analyzer RNA QC (Agilent). RNAseq libraries were constructed by the Transcriptional Regulation and Expression (TREx) Facility at Cornell University using the NEBNext Ultra II RNA Library Prep Kit (New England Biolabs).

## Notes

This work was supported by R01AI139664 (CL) and HHMI Gilliam Fellowship (KGM).

### Competing Interest Statement

The authors have declared no competing interest.

