## Supplemental Information for "Stimulator of interferon genes is required for Toll-Like Receptor-8 induced interferon response"

**Supporting information:**

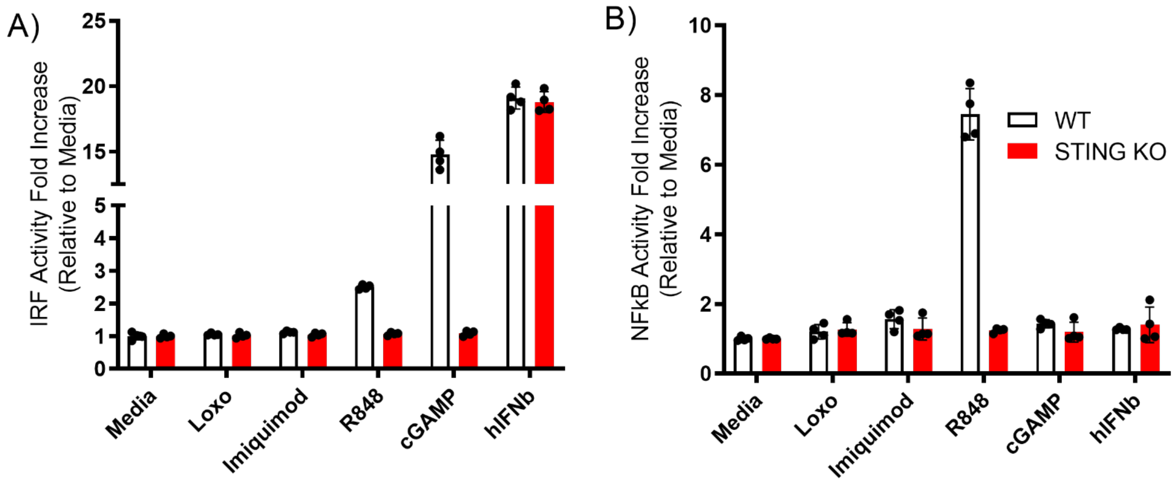


**Supplementary Figure 1**. WT and STING KO DTHP-1 cells were stimulated with 10 µg/mL Loxoribine, 10 µg/mL imiquimod, 10 µg/mL R848, , 10 µg/mL cGAMP, or 100U/mL IFNβ for 18 hours. Supernatants were collected and A) IRF activity was measured using Lucia Luciferase activity n=3. B) NFkβ activity was measured using SEAP activity, n=3.

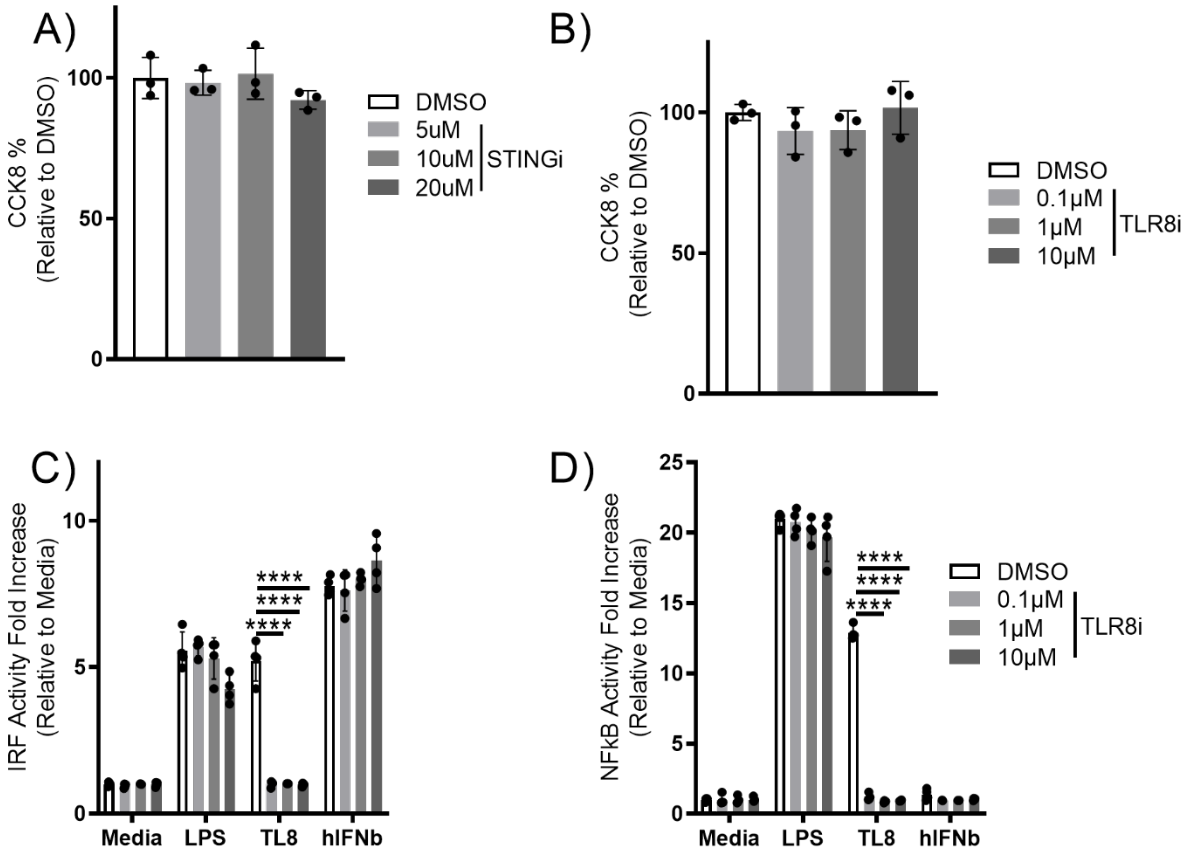


**Supplemental Figure 2. SN0-11 and CUCPT9a inhibitors are not toxic to DTHP-1 cells.** DTHP-1 cells were treated with vehicle or increasing concentrations of SN-011 for 18 hours, then CCK8 reagent was added and CCK8 activity was measured after 2 hours. A) CCK8 activity in SN0-11 treated DTHP-1 cells, n=3. DTHP-1 cells were treated with vehicle, or increasing concentrations of CUCPT9a for 18 hours, then CCK8 reagent was added and CCK8 activity was measured after 2 hours. B) CCK8 activity in CUCPT9a treated cells, n=3. DTHP-1 cells were inhibited with increasing concentrations of CUCPT9a for 1hr, then stimulated with 1μg/mL LPS or 10 μg/mL TL8 for 18 hours. C) IRF activity was measured using Lucia Luciferase activity. D) NFkβ activity was measured using Secreted Alkaline Phosphatase activity, n=3.

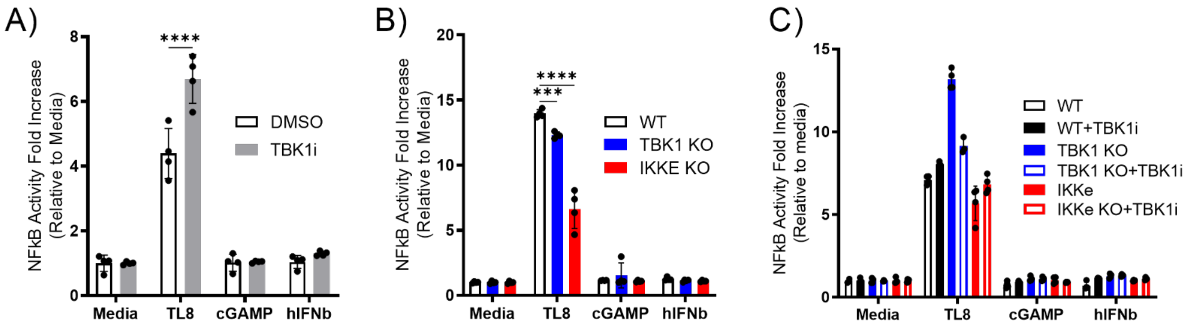


**Supplemental Figure 3: TL8 induces STING phosphorylation and requires IKKε, but not TBK1.** WT DTHP-1 cells were treated with vehicle, 10µM MRT67307 for 1hr then stimulated with TL8, cGAMP, or hIFNβ for 18 hours. A) NFkβ activity was measured using SEAP activity, n=6. WT, TBK1 KO, or IKKε KO DTHP-1 cells were stimulated with TL8, cGAMP, or hIFNβ for 18 hours. B) NFkβ activity was measured using SEAP activity, n=6. WT, TBK1 KO, or IKKε KO DTHP-1 cells were treated with vehicle or 10µM MRT67307 for 1hr then stimulated with TL8, cGAMP, or hIFNβ for 18 hours. C) NFkβ activity was measured using SEAP activity, n=3.

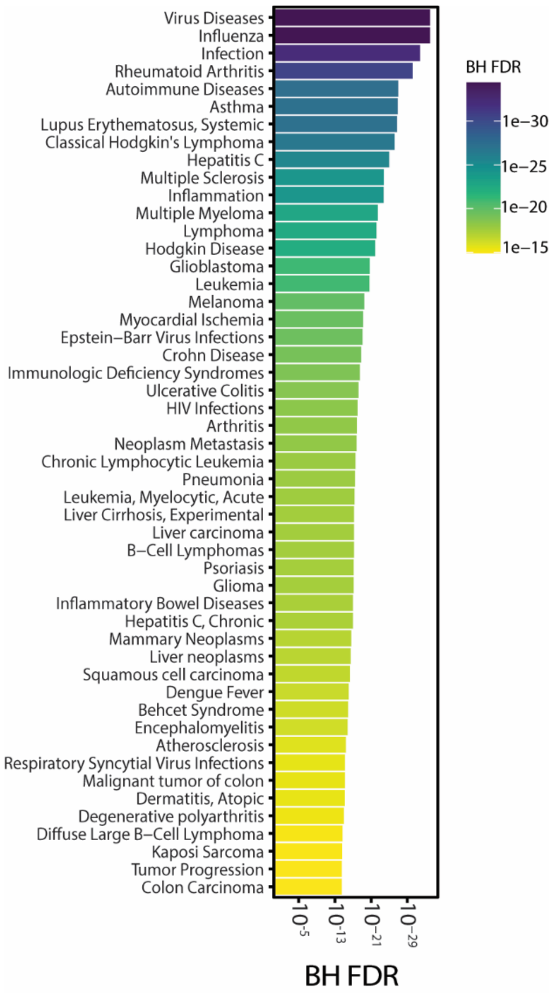


**Supplemental Figure 4. TL8 stimulation induces genes associated with infection, cancer, and autoimmune disease.**

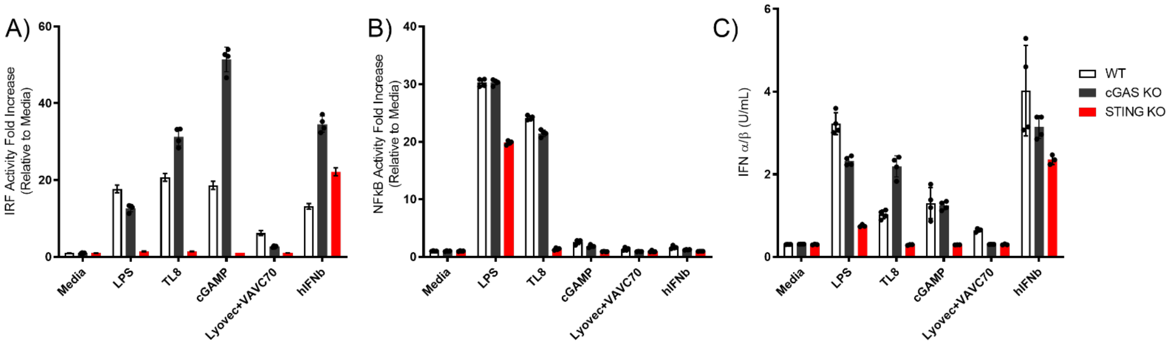


**Supplementary Figure 5**. WT, cGAS KO, and STING KO DTHP-1 cells were stimulated with), 1 μg/mL LPS, 10 μg/mL TL8, 10 μg/mL cGAMP, 10 μg/mL Lyovec+VAVC70, or 100U/mL IFNβ for 18 hours. Supernatants were collected and A) IRF activity was measured using Lucia Luciferase activity, n=3. B) NFkβ activity was measured using SEAP activity, n=3. C) IFNα/β secretion was measured using IFNα/β Reporter HEk293 cells, n=3.
